# Evaluation of Significance of *SERPINA3* and *SPP1* in Driving Glioma Progression Through Dysregulation of Brain Inflammatory Pathway

**DOI:** 10.64898/2026.02.11.705371

**Authors:** Hirtik Singh Rathore, Siya Singh, Sukhmanpreet Singh, Jassi Goyal, Dibyajyoti Banerjee

## Abstract

Glioma is a highly aggressive malignancy with a poor prognosis, particularly in grade IV glioblastoma. The monitoring of disease progression remains challenging due to high heterogeneity in the tumour and a lack of progressive markers to keep track of the tumour progression. This scarcity of good progressive biomarkers has led us to search for better options. SERPINA3 and SPP1 were investigated as potential dual biomarkers reflecting tumour microenvironment and enabling assessment of progression. In silico analysis was conducted, where correlation analysis was performed to evaluate the cell-type specificity of SERPINA3 in astrocytes and SPP1 in microglial cells, with comparisons across other neural populations in both low-grade glioma and glioblastoma. Pan-cancer expression analysis was conducted to determine whether these biomarkers remain significantly elevated in glioma relative to other malignancies, including hepatocellular carcinoma, given their hepatic origin. Alzheimer’s disease datasets were analysed to verify the relevance in neurodegenerative diseases. The in-silico analysis revealed that SERPINA3 and SPP1 show astrocytic and microglial specificity, respectively, and exhibit their highest expression levels in glioma across cancers. Co-expression analysis further identified enrichment of immunoregulatory pathways alongside upregulated oxidative stress–associated markers, highlighting the functional relevance of these biomarkers within the glioma microenvironment.

## Introduction

The survival time of GBM patients is only 12–15 months, making it the most lethal type of brain tumour. Every year, about 200,000 people worldwide succumb to this disease[1,2]. These tumours arise from neural stem or progenitor cells that acquire tumour-initiating genetic alterations and grow as highly infiltrative lesions within the brain parenchyma. According to the World Health Organisation (WHO) classification of CNS tumours, gliomas are graded from I to IV based on histopathological and molecular characteristics. Grade I gliomas are generally biologically benign with low risk, whereas Grade II gliomas, show a substantial tendency for recurrence. Grade III (anaplastic gliomas) and Grade IV gliomas, known as glioblastoma multiforme (GBM), are high-grade, poorly differentiated, and malignant tumours associated with an extremely poor prognosis[3]. Glioblastoma malignancies occur as diffuse tumours that infiltrate the brain parenchyma, contributing to poor survival and inevitable recurrence[4].

A major challenge in glioma management is the diffuse and heterogeneous nature of these tumours. Due to pronounced intra-tumoral heterogeneity, a biomarker measured from a single biopsy may not represent the entire tumour biology, leading to inaccurate prognostic predictions[4]. This heterogeneity is a major obstacle to reliable disease monitoring and limits the establishment of biomarkers that are sensitive, specific, and accessible through non-invasive or minimally invasive methods. As a result, only a few biomarkers, such as isocitrate dehydrogenase (IDH) mutations and O^6^-methylguanine-DNA methyltransferase (MGMT) promoter methylation, have consistently demonstrated prognostic or predictive value in gliomas[5].

In glioblastoma and other gliomas, accurate assessment of tumour progression and aggressiveness is essential for prognosis and monitoring response to therapy, so that the appropriate treatment can be given in accordance to the progression of the disease. However, currently used biomarkers do not sufficiently capture the complex interactions between tumour cells and the surrounding microenvironment, which are essential for glioma progression.

Biomarkers that reflect tumour–microenvironment interactions are of particular interest. Secreted phosphoprotein 1 (SPP1) and serpin family A member 3 (SERPINA3) represent such candidates, as they capture immune contributions that are critical drivers of glioma progression. Serpins constitute a large family of proteins primarily involved in the regulation of serine protease activity. Among them, SERPINA3 (also known as α1-antichymotrypsin) is one of the most extensively characterised in human pathology and is predominantly produced by the liver and astrocytes [6, 7]. In cancer cells, the nuclear fraction of SERPINA3 activates the MAPK/ERK1/2 and PI3Kδ signalling pathways through stimulation of NF-κB signalling and AKT phosphorylation, resulting in inhibition of the G2/M phase transition, anti-apoptotic effects, and enhanced tumour proliferation[8]. Overexpression of SERPINA3 is positively correlated with glioma development and poor prognosis, and it promotes remodelling of astroglia and the microglial extracellular matrix[7–9].

*SPP1*, also known as osteopontin, is a phosphorylated glycoprotein with major roles in cell adhesion and migration, as well as signalling pathways. Recent evidence shows high expression of *SPP1* in a variety of tumours such as colorectal cancer, breast cancer, and more. Models explaining the role of osteopontin include downstream activation of the NF-κβ signalling pathway. SPP1 contributes to cancer progression by promoting tumour growth. *SPP1* is overexpressed in glioma, particularly in high-grade glioblastomas. Acts as a pro-tumorigenic cytokine, interacting with integrins and CD44 on glioma cells to promote migration and survival.[10] High SPP1 expression correlates with poor prognosis. Due to its role in immune evasion, SPP1 is being explored as a target for glioma therapy[11]. The present study aimed to explore the relationship of *SERPINA3 and SPP1* expression with respect to glioma using multiomics data of lower-grade gliomas and glioblastoma. To further understand this relationship a correlation analysis was done, and a gene co-expression analysis was also done to find out the upregulated pathways that are involved with these biomarkers, giving us an insight into the tumour microenvironment.

## Materials and methods

This study was conducted as a computational (in silico) analysis using transcriptomic, proteomic, and clinical datasets to evaluate *SERPINA3* and *SPP1* as glioma progression biomarkers. Gene expression data from primary human tumours, including low-grade glioma (LGG) and glioblastoma (GBM), were taken from OncoDB (https://oncodb.org). The Cancer Genome Atlas (TCGA) was accessed through the Genomic Data Commons portal (https://portal.gdc.cancer.gov). Pan-cancer expression profiling was performed using the UALCAN platform (https://ualcan.path.uab.edu), which provides access to TCGA-derived normalised transcriptomic data and associated clinical annotations.

To check disease specificity, transcriptomic data from hepatitis were obtained from the Gene Expression Omnibus (GEO) dataset GSE143318 (https://www.ncbi.nlm.nih.gov/geo/) as both biomarkers are of hepatic origin. Gene expression data from normal human brain tissue were retrieved from the Genotype-Tissue Expression (GTEx) project (https://gtexportal.org). Disease-specific datasets, including Alzheimer’s disease transcriptomic profiles, were accessed from the AlzData platform (http://www.alzdata.org).

### Cell-Type–Specific Marker Selection

Both SERPINA3 and SPP1 are upregulated in glioma, but also by reactive astrocytes and tumour-associated microglia as part of the tumour microenvironment. To check cell specificity, molecular markers were selected for each type of cell populations. *GFAP, ALDH1L1* are markers for astrocytes[12]. *HEXB* [13] and *TMEM119* [13–15]markers for microglia. *BMP4, ENPP4, and ASPA* markers for oligodendrocytes [11]. *MAP2, SYN1*, and *TUBB3* for neuronal markers [16, 17]These markers were used to check the association of *SERPINA3* and *SPP1* with astrocytes and glial cells.

### Correlation and Co-expression Analysis

Correlation analyses using Spearman and Pearson’s methods were performed between *SERPINA3* and *SPP1* expression levels and the cell-specific markers in both LGG and GBM. Co-expression profiling was conducted to identify the top 50 genes positively correlated with *SERPINA3* and *SPP1*. These analyses were used to determine which cell type the co-expressed genes were enriched. High co-expression in specific cells indicates potential roles in glioma progression.

### Differential Expression and Pan-Cancer Analysis

Differential gene expression analysis of *SERPINA3* and *SPP1* was performed in LGG and GBM, and pan-cancer data analysis was done using the UALCAN platform (https://ualcan.path.uab.edu). Log-2-fold change values were calculated, and the p-values were used to check statistical significance. Proteomic expression data for these biomarkers in GBM were obtained from OncoDB to check if it correspondes with transcriptomic findings.

### Alzheimer’s Disease Comparison

To confirm that the observed upregulation of *SERPINA3* and *SPP1* is glioma-specific and not a feature of neuroinflammatory diseases, transcriptomic datasets from Alzheimer’s disease (AD) patients were analysed. Expression patterns across key brain regions (hippocampus, entorhinal cortex, frontal and temporal cortices) were compared to glioma samples to check disease specificity.

### Proliferation Marker Association

To evaluate the association between these biomarkers and tumour proliferation, correlation analyses were performed with cell proliferation markers *MKI67, PCNA and TOP2A*. Markers of cell proliferation, *MKI67 (Ki-67)*[18], *PCNA* [19], *and TOP2A*[20], are commonly used as an indicator of glioblastoma cells dividing.

Studies using immunohistochemistry have shown that *TOP2A and PCNA* levels tend to rise together in gliomas, reflecting high proliferative activity [19]and *TOP2A* expression also correlates with *Ki-67* levels in GBM samples[20]. Together, it highlights the reason for examining multiple proliferation markers to understand tumour growth in glioblastoma. These analyses were conducted in GBM datasets to assess whether the selected marker levels relate to the increased proliferation.

### Statistical Analysis and Data Visualisation

Correlation and co-expression analyses were performed using the R Studio (version 4). Gene expression matrices of the biomarkers and selected cell-type–specific markers were taken from the LGG and GBM datasets. Correlations between both biomarkers and cell-type markers were made using Spearman and Pearson correlation coefficients, and statistical significance was adjusted to p < 0.05 using multiple testing using the Benjamini-Hochberg method. Co-expression profiling was conducted to identify the top 50 genes positively correlated with *SERPINA3* and *SPP1*. The corrplot and pheatmap packages were used to generate heatmaps of co-expression patterns. High correlation with cell-type–specific markers was interpreted as an indication of cell-specific expression, thereby linking *SERPINA3* and *SPP1* expression to particular glial or neuronal lineages.

## Results

### Transcriptomic and Proteomic analysis of SERPINA3 and SPP1 in GBM and LGG

To evaluate the elevation of expression of SERPINA3 and SPP1 in glioma grades mRNA expression data were extracted from oncoDB, and boxplot comparisons were done to evaluate differential expression between normal brain tissue and glioma samples in low-grade glioma (LGG) and glioblastoma (GBM).As demonstrated in ***Fig.1*** *SERPINA3* expression was elevated in both LGG and GBM (Student’s t-test, p-values < 0.05) compared to normal tissues, which showed minimal expression. Its expression increased in GBM relative to LGG, indicating tumour progression. A similar pattern was observed for *SPP1*. Significantly higher *SPP1* expression was present in LGG and was even more increased in GBM (Student’s t-test, p-values < 0.05) when compared with normal tissue samples. While both glioma grades show upregulation relative to the normal brain, the magnitude and statistical significance of *SPP1* expression were greater in GBM.

**Fig 1.**
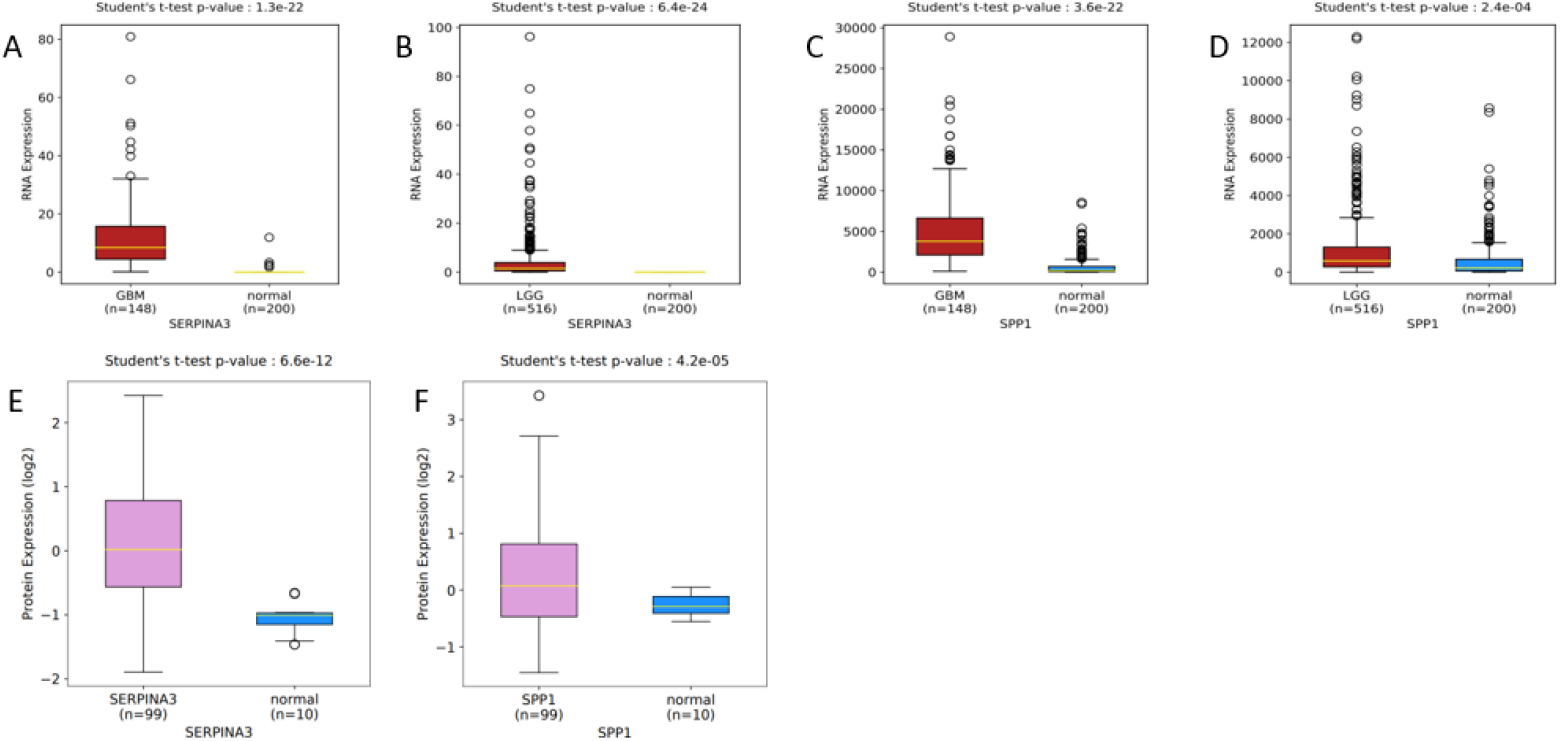
Transcriptomic and proteomic analysis of SERPINA3 and SPP1 expression in glioblastoma multiforme (GBM) and low-grade glioma (LGG). **(A)** SERPINA3 RNA expression in GBM (n = 148) compared with normal brain tissue (n = 200). **(B)** SERPINA3 RNA expression in LGG (n = 516) compared with normal brain tissue (n = 200). **(C)** SPP1 RNA expression in GBM (n = 148) compared with normal brain tissue (n = 200). **(D)** SPP1 RNA expression in LGG (n = 516) compared with normal brain tissue (n = 200). **(E)** SERPINA3 protein expression in GBM compared with normal brain tissue. **(F)** SPP1 protein expression in GBM samples (n = 99) compared with normal brain tissue (n = 10). Statistical significance was defined as *p* < 0.05.

Proteomic abundance of *SERPINA3* and *SPP1* was checked in GBM samples and compared with normal controls to determine whether the transcriptomic levels correspond to increased protein expression. Measuring protein levels provides a complete picture of efficient translation. In ***Fig.1(E), (F)*** protein-level analyses confirmed that the elevated mRNA expression corresponds into increased protein levels, indicating involvement of these molecules in the tumour microenvironment. The *p*-value <0.05 for both markers indicates that this difference is highly reliable and prominent.

### To verify astrocytic localization of SERPINA3 and SPP1

To evaluate the localisation of *SERPINA3* in astrocytes, log2-normalised RNA expression levels of astrocyte markers *GFAP* and *ALDH1L* were correlated with *SERPINA3* using Pearson’s method and Spearman’s method in GBM and LGG cohorts. In ***Fig.2*** Correlation coefficient r = 0.472 for *GFAP* and r =0.34 for *ALDH1L*, indicating a moderately positive correlation, which is highly significant (p-values < 0.05) in GBM. High and low groups of *GFAP* and *ALDHL1* are stratified and compared with *SERPINA3* expression levels using Student’s t-test. Gliomas with high *GFAP* and *ALDHL1* expression had significantly elevated *SERPINA3* levels. Hence, *SERPINA3* is associated with astrocytic differentiation and reactive glioma. A similar pattern was observed for Low-grade Glioma, where *SIRPINA3* was correlated with the astrocyte biomarkers *ALDH1L1* and *GFAP. SPP1* shows a weak to moderate association with astrocytic markers, hinting at active communication between microglia and astrocytes.

**Fig 2.**
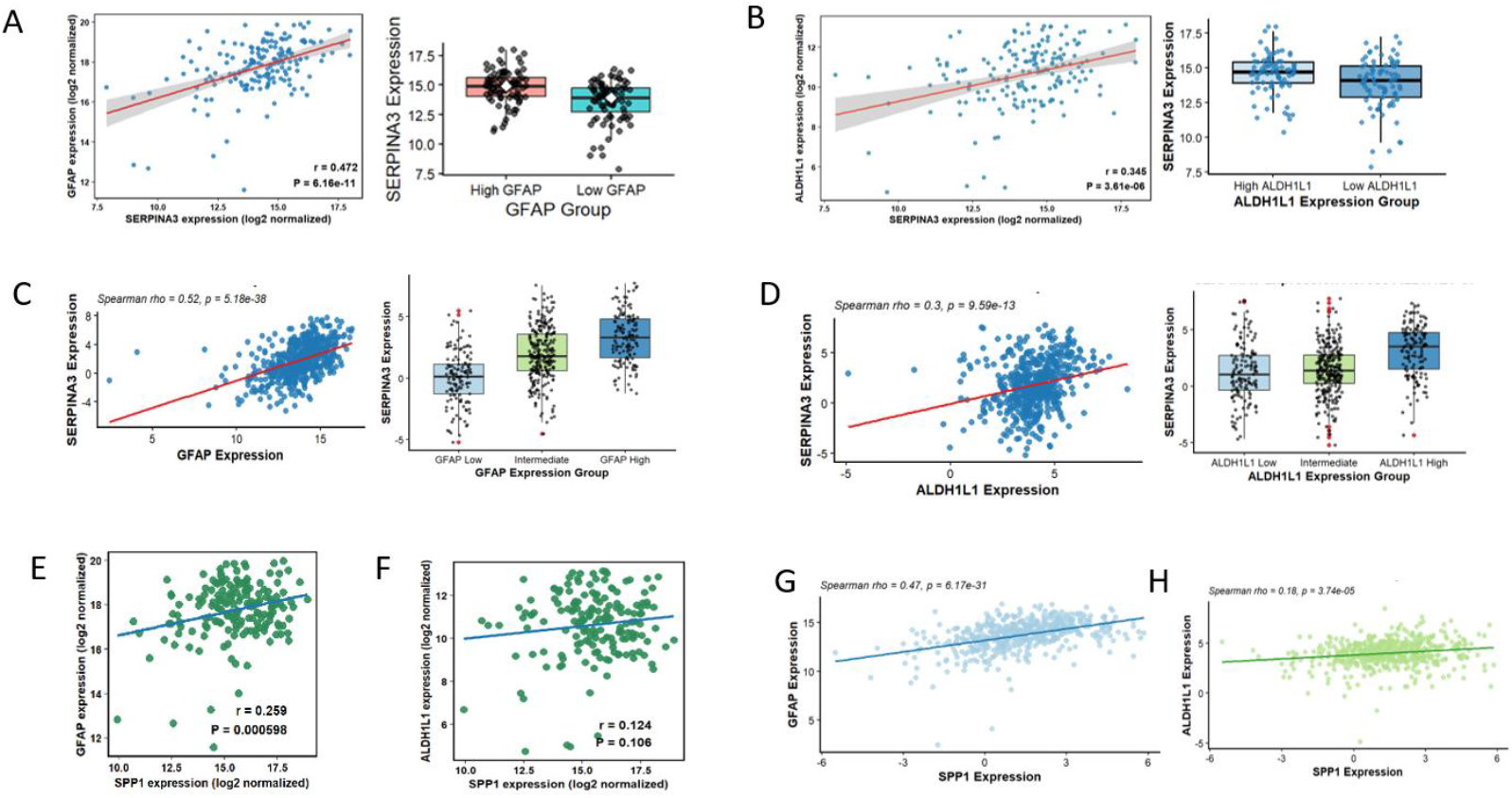
Localisation of SERPINA3 in astrocytes in glioblastoma. Transcriptomic data of GBM and LGG from the TCGA-GBM cohort were analysed using log_2_-normalized RNA expression values. **(A)** SERPINA3 showed a positive correlation with GFAP expression (*r* = 0.472, *p* = 0.05), with elevated SERPINA3 levels observed in the GFAP-high group relative to the GFAP-low group in GBM. **(B)** A significant positive correlation between SERPINA3 and ALDH1L1 expression (*r* = 0.345, *p* >0.05) was observed in GBM **(C)** correlation with GFAP is strong and highly significant (ρ = 0.52, p < 0.05) in LGG **(D)** SERPINA3 shows a moderate positive correlation with the astrocytic marker ALDH1L1 (ρ = 0.3, p < 0.05) in LGG (E) SPP1 exhibits a weak positive correlation with the astrocytic marker GFAP (r = 0.259, p < 0.05) in GBM, indicating astrocytic contribution. **(F)** The correlation of SPP1 with ALDH1L1 is very weak and not significant (r = 0.124, p > 0.05) in GBM. **(G)** SPP1 shows a moderate positive correlation with the GFAP (ρ = 0.47, p< 0.05), indicating astrocytic association correlation in LGG. **(H)** SPP1 with ALDH1L1 is weak but significant (ρ = 0.18, p < 0.05) in LGG. Pearson correlation analyses in **(A), (B), (E)** and **(F)** were performed for GBM. Spearman correlation analyses in **(C), (D), (G)** and **(H)** were performed for LGG.

### To verify microglial, oligodendrocytes and neuronal specific expression pattern of SERPINA3 and SPP1

We observed a general trend in ***Fig.4*** where higher *SERPINA3* expression was linked with lower expression of oligodendrocyte markers. Although weak positive correlations were detected, their very small value has limited biological relevance. The observed negative correlations are statistically insignificant and therefore should be interpreted as indicative of a trend in oligodendrocytes. A similar non-significant negative trend was observed between *SERPINA3* and neuronal markers, suggesting a possible reduction in neuronal integrity in inflammation-rich tumour environments. With respect to microglial markers in ***Fig.3*** *SERPINA3* a stronger correlation only with *HEX-B*, while other microglial markers exhibited weak or negligible correlations. According to a previous study, overexpression of *SERPINA3* in astrocyte/microglia co-cultured GSCs was observed[9], which may be indicative of astrocyte microglial cell crosstalk that may be occurring here as well. In ***Fig.5*** Neuronal cells showcased a negative correlation, but can be considered as a general trend or there can be caused by neural degeneration due to glioma. The results validate the specificity of *SPERNA3* in astrocytes and *SPP1* in microglial cells and together, these findings point to an innate immune–driven glial interaction.

**Fig 3.**
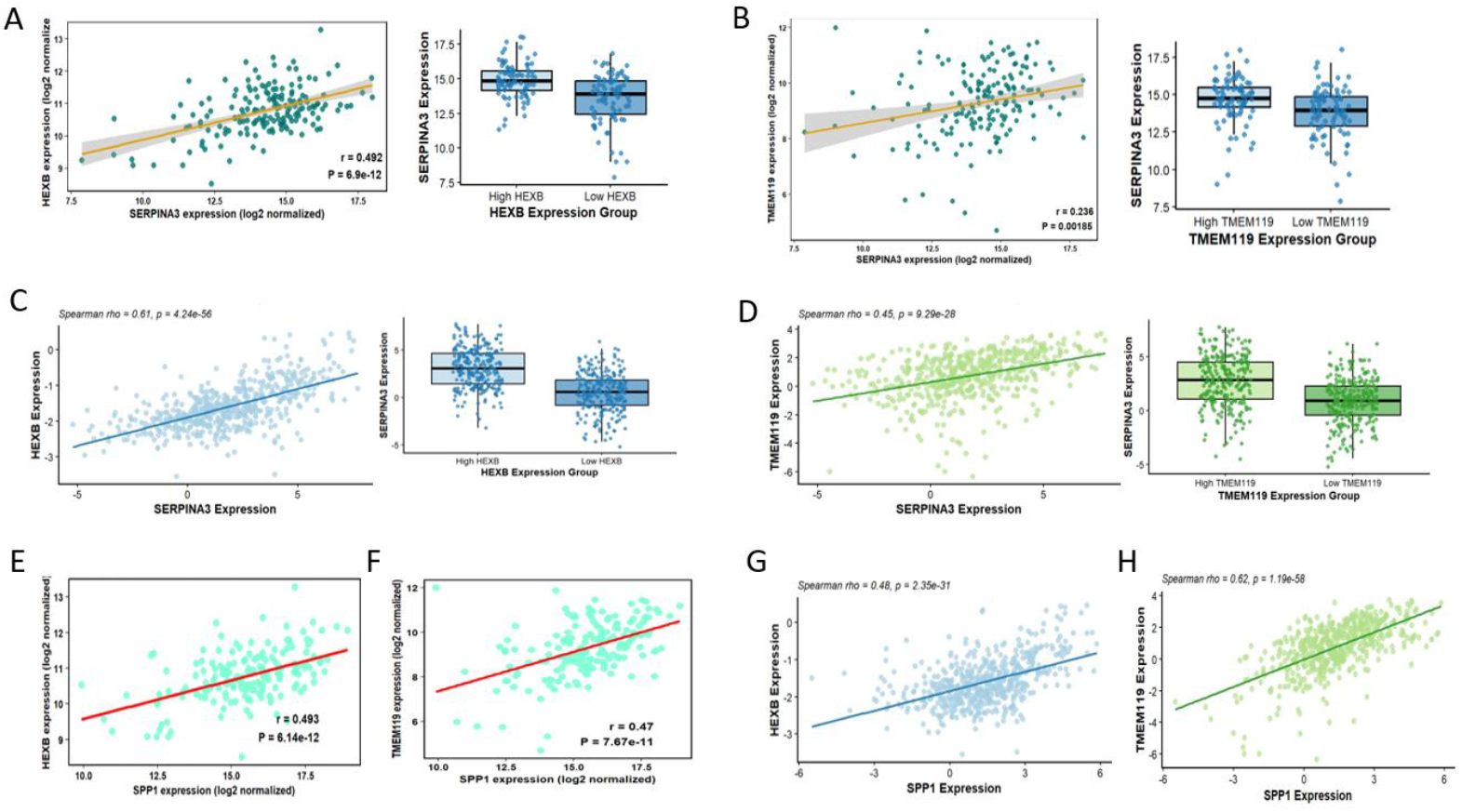
Correlation analysis using transcriptomic data from GBM and LGG samples, highlighting microglia-specific expression patterns. **(A)** shows a strong positive correlation between SERPINA3 and the microglial marker HEXB (*r* = 0.492, *p*<0.0.5), with higher SERPINA3 expression in HEXB-high samples compared with HEXB-low samples in GBM. **(B)** weakly positive correlation between SERPINA3 and microglial marker, TMEM119 (*r* = 0.236, *p* <0.05) supported by increased SERPINA3 expression in TMEM119-high samples in GBM **(C), (D)** In LGG, SERPINA3 exhibits strong positive correlations with HEXB (ρ = 0.61, p < 0.05) and TMEM119 (ρ = 0.45, p < 0.05), indicating dominant microglial linkage. **(E) (F)** In GBM, SPP1 shows strong positive correlations with HEXB (r = 0.493, p < 0.05) and TMEM119 (r = 0.47, p < 0.05), reinforcing microglial specificity. **(G) (H)** In LGG, SPP1 correlates positively with HEXB (ρ = 0.48, p < 0.05) and strongly with TMEM119 (ρ = 0.62, p < 0.05). Pearson correlation analyses in **(A) (B), (E)** and **(F)** were performed for GBM. Spearman correlation analyses in **(C), (D), (G)** and **(H)** were performed for LGG.

**Fig 4.**
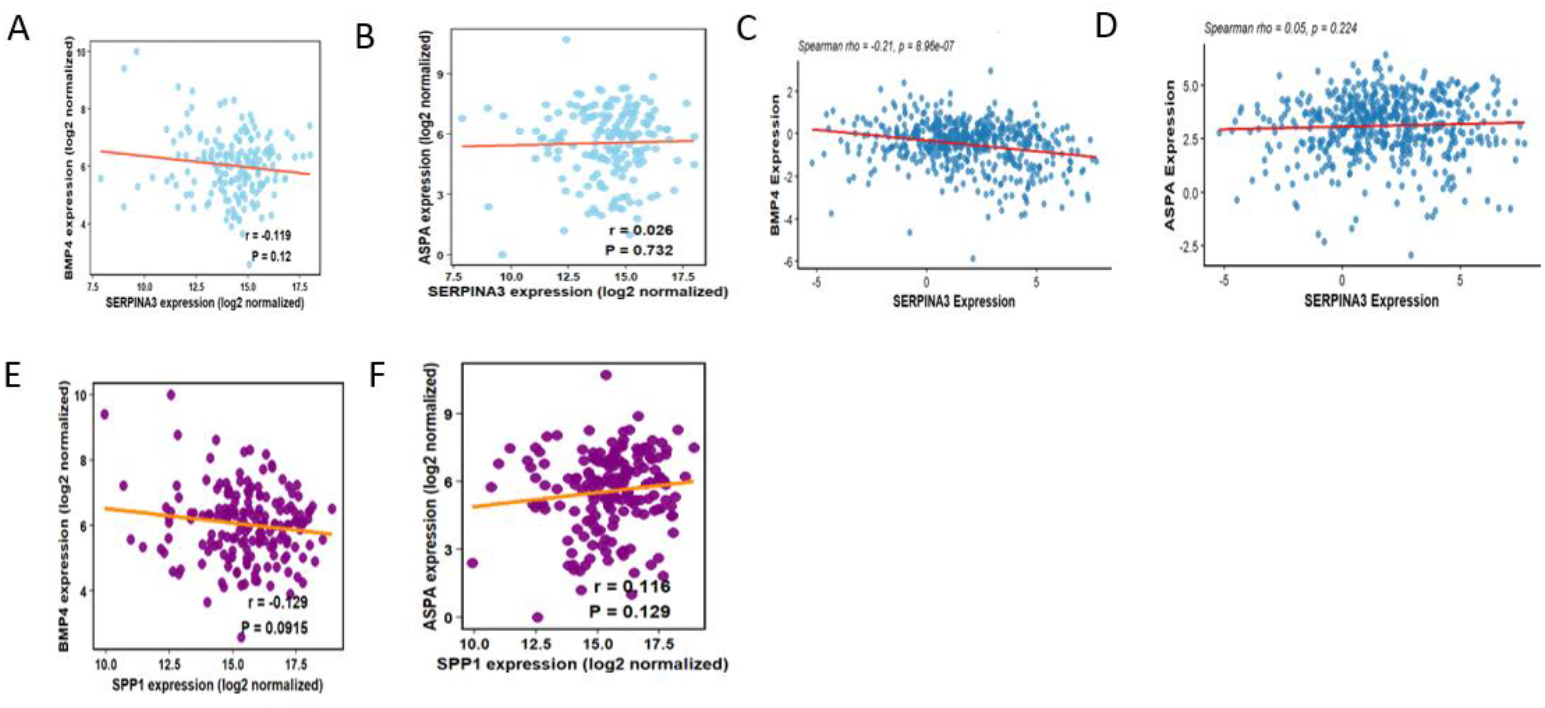
Correlation analysis using transcriptomic data from GBM and LGG samples, highlighting oligodendrocyte-specific expression patterns **(A), (B)** show correlations between SERPINA3 and oligodendrocyte lineage markers BMP4 (*r* = −0.119, *p* >0.0.5), ASPA (*r* = 0.026, *p* > 0.05) in GBM **(C)** The correlation with the oligodendrocyte marker BMP4 is very weakly negative and not significant (r = −0.129, p < 0.05) negligible oligodendrocyte involvement in LGG. **(D)** a very weak positive correlation with ASPA (ρ = 0.05, p > 0.05), indicating minimal oligodendrocyte involvement in LGG. **(E)** The weakly negative correlation with the oligodendrocyte marker BMP4 and non-significant (r = −0.129, p > 0.05), indicating negligible oligodendrocyte involvement in GBM. **(F)** SPP1 shows a very weak positive, non-significant correlation with ASPA (r = 0.116, p > 0.05), consistent with minimal oligodendrocyte association in GBM. Pearson correlation analyses in **(A) (B), (E)** and **(F)** were performed for GBM. Spearman correlation analyses in **(C)** and **(D)** were performed for LGG.

**Fig 5.**
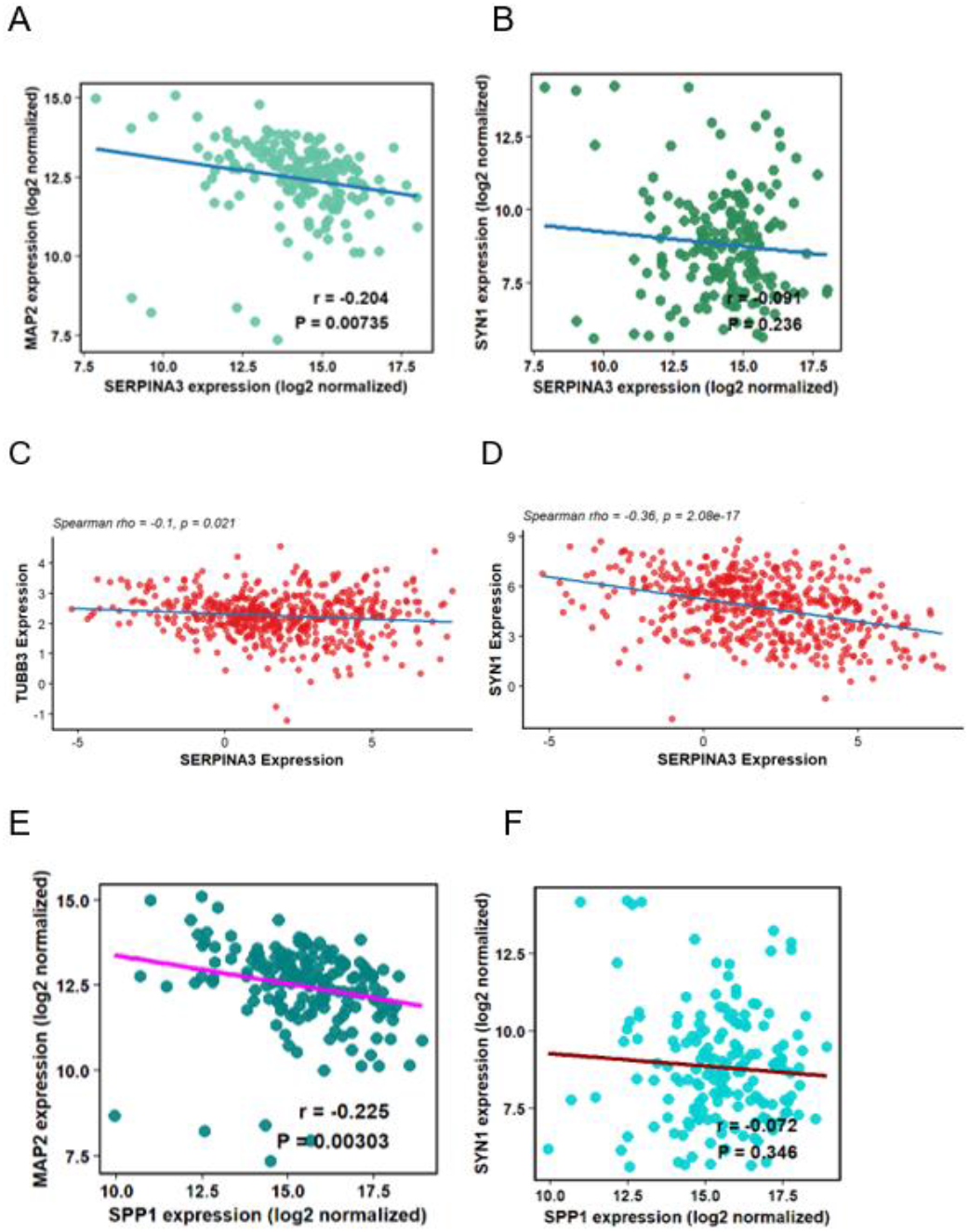
Correlation analysis using transcriptomic data from GBM and LGG samples, highlighting neuronal-specific expression patterns **(A)** and **(B)** show minimal correlations between SERPINA3 and neuronal markers MAP2 (*r* = −0.204, *p* >0.05) and SYN1 (*r* = −0.090, *p* >0.05), revealing weak negative correlations which are not significant in GBM **(C) (D)** SERPINA3 shows a weak-to-moderate negative correlation with neuronal markers, including TUBB3 (ρ = −0.1, p < 0.05) and SYN1 (ρ = −0.36, p < 0.05), suggesting negative association with neuronal identity in LGG **(E)** In contrast, SPP1 shows a weak-to-moderate significant negative correlation with MAP2 (r = −0.225, p < 0.05), suggesting an inverse relationship with neuronal identity in GBM. **(F)** The correlation with the neuronal marker SYN1 is very weakly negative and not significant (r = −0.072, p > 0.05), indicating lack of neuronal coupling in GBM. Pearson correlation analyses in **(A) (B), (E)** and **(F)** were performed for GBM. Spearman correlation analyses in **(C)** and **(D)** were performed for LGG.

### Pan cancer expression analysis by UALCLAN

To further evaluate the specificity of *SERPINA3* and *SPP1* as glioma biomarkers, their expression was assessed across multiple cancer types using pan-cancer transcriptomic datasets as *SPP1* and *SERPINA3* are inflammatory and tumour-associated proteins that are not restricted to a single tissue. They are known to be predominantly produced in the liver, in addition to being associated with inflammatory and proliferative processes in other tissues. Because of this hepatic origin, their levels are expected to be elevated in hepatocellular carcinoma (HCC)[10, 21, 22].This analysis aimed to determine whether the upregulation of these genes is unique to gliomas or represents a broader oncogenic pattern observed in other malignancies. In ***Fig.6*** *SERPINA3* is upregulated across multiple cancer types, including GBM, LIHC, LUSC, PAAD, and COAD. A similar pattern was observed in *SPP1*, where various cancer types were showcasing a rising expression of *SPP1* raising questions regarding their tumour specificity. Despite this, *SERPINA3* and *SPP1* exhibit highest expression levels in GBM. We shall take into account that the expression of *SERPINA3* in other cancers is a result of peripheral expression of this inflammatory biomarker, which cannot cross the blood-brain barrier; hence, both can be considered specific when it is correlated within the context of CSF. Therefore, we can consider it as a progressive biomarker.

**Fig 6.**
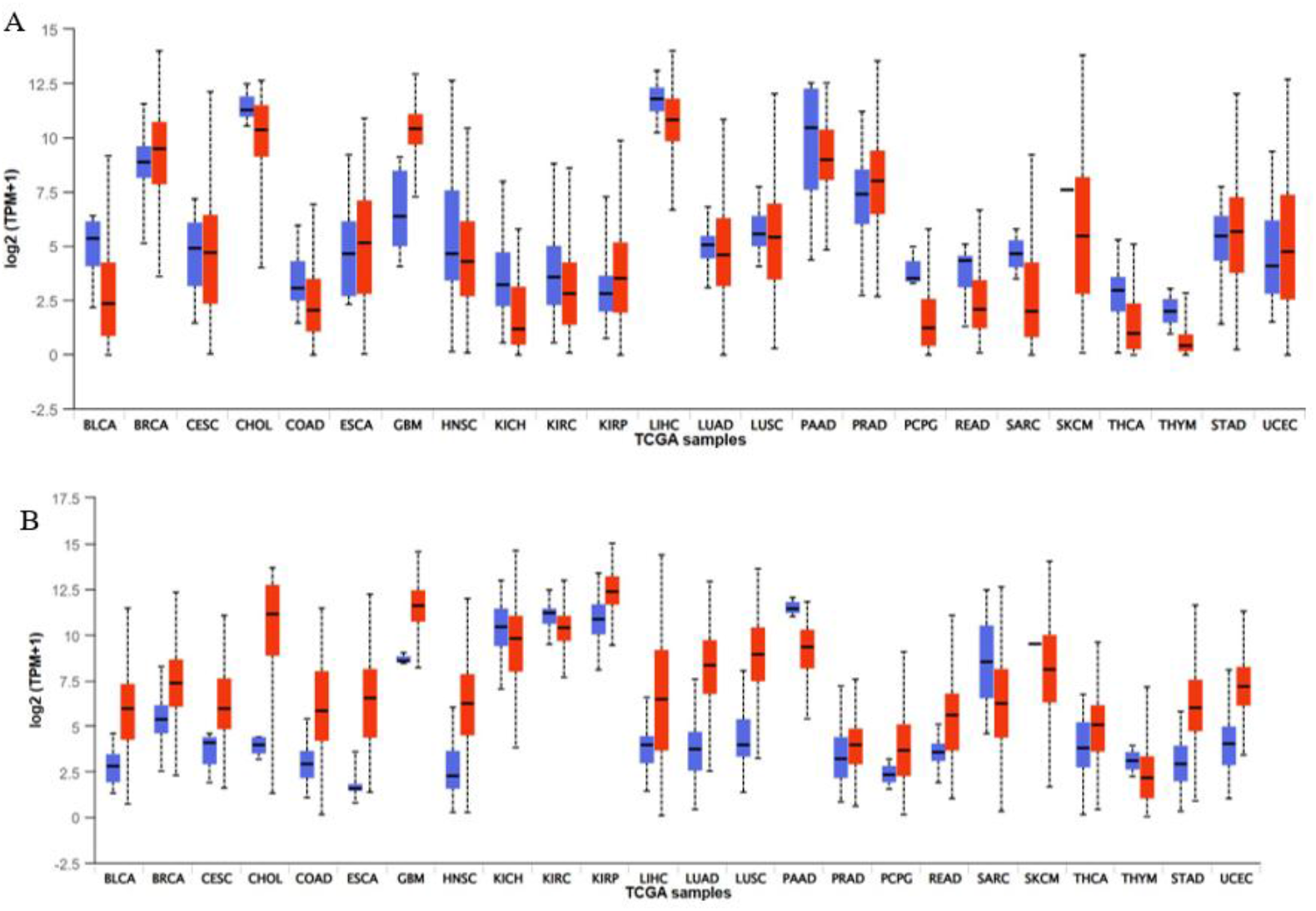
Pan-cancer expression analysis of SPP1 and SERPINA3 across TCGA cohorts. RNA sequencing data from The Cancer Genome Atlas (TCGA) were analysed to evaluate the expression patterns of *SPP1* and *SERPINA3* across multiple cancer types using tumour and corresponding normal tissue samples, with expression values represented as log_2_(TPM + 1) **(A)** The pan-cancer expression profile of *SPP1*, showing significant tumour-associated upregulation in several cancer types, it is highest in glioma cohorts, relative to normal tissues. **(B)** The expression pattern of *SERPINA3* across TCGA cancers demonstrates elevated levels in selected tumour tissues compared with matched normal controls

### SERPINA3 and SPP1 levels in Alzheimer’s disease show no correlation with the disease

*SERPINA3* [23–25]and *SPP1* [26–28] have been reported to show elevated expression in certain neurodegenerative conditions, including Alzheimer’s disease. To ensure that their association is specific to glioma, their expression was checked in transcriptomic datasets from Alzheimer’s patients. In ***Fig.7 (A) and (B)*** *SERPINA3* elevation is most present in the temporal cortex. *SPP1* upregulation is strongest in the frontal cortex. The pathology of Alzheimer’s disease originates within the hippocampus and entorhinal regions, and subsequently spreads throughout the frontotemporal cortices. It reaches as far as the striatum and thalamus, usually with sparing of the cerebellum[29] If they were to be Alzheimer’s inflammatory marker, one would expect major and even more pronounced changes in the entorhinal cortex, hippocampus as well. Both markers fail to show strong significance in these regions so it cannot be called as an AD marker.

**Fig 7.**
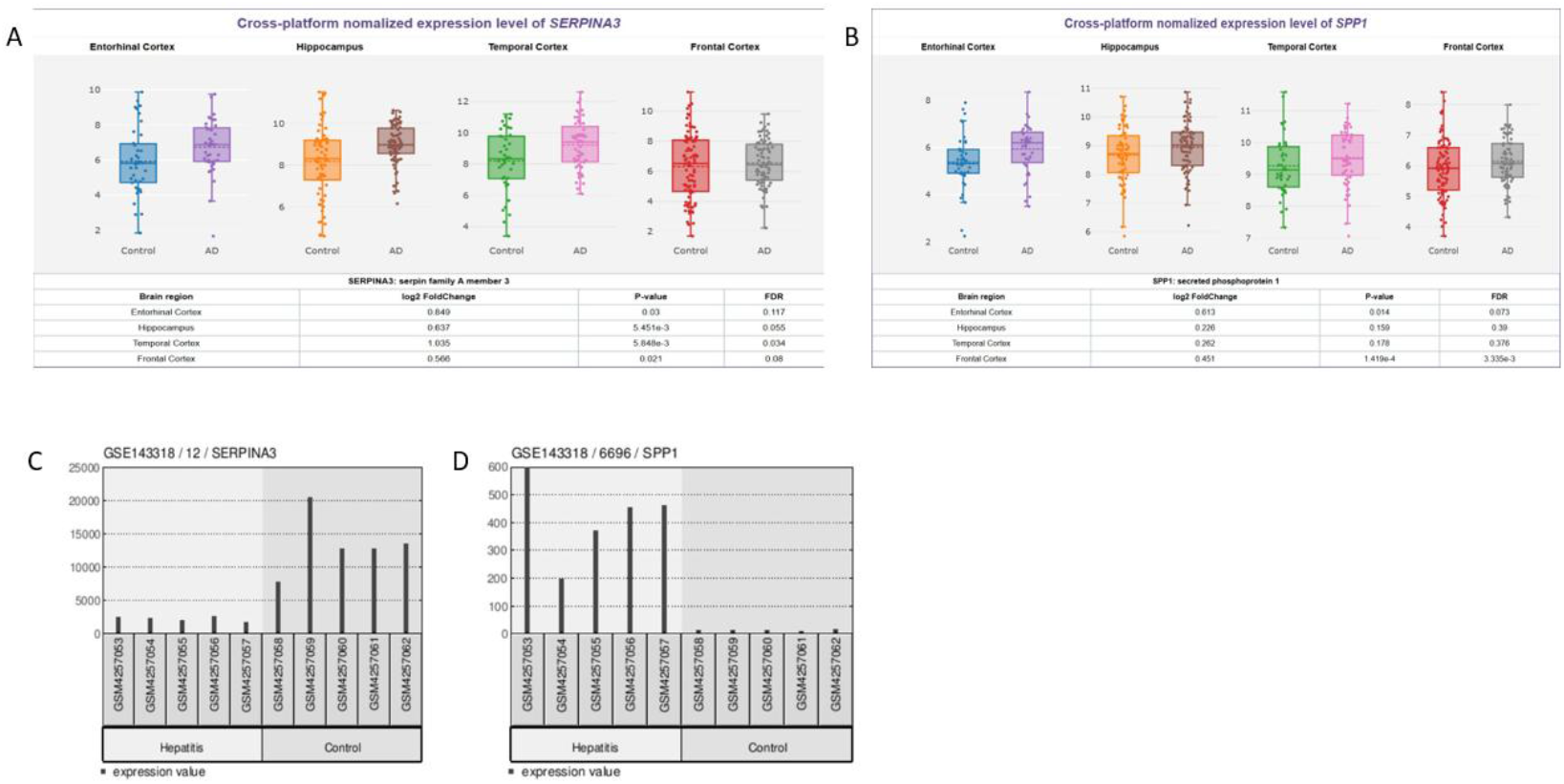
Normalised expressions of SERPINA3 and SPP1 to investigate the relation of these biomarkers with Alzheimer’s disease and hepatitis. **(A)** illustrates the cross-platform normalised expression levels of SERPINA3 across the entorhinal cortex, hippocampus, temporal cortex, and frontal cortex in control and Alzheimer’s disease (AD) samples. SERPINA3, the highest biologically relevant log2 fold change observed in the temporal cortex, followed by the hippocampus **(B)**, depicts the expression profile of SPP1 across the same brain regions, where statistically significant differences are primarily confined to the entorhinal cortex and frontal cortex, while changes in the hippocampus and temporal cortex are biologically less pronounced. **(C)** shows individual sample expression of SERPINA3 in hepatitis and **(D)** SPP1 shows elevated expression across the same samples, with each bar representing a single GEO sample. Comparisons between hepatitis and control groups were performed, confirming that only SPP1 is upregulated in hepatitis samples relative to controls.

### SERPINA3 and SPP1 comparison with hepatitis

*SERPINA3* and *SPP1* are known to be actively expressed in the liver. To assess the disease specificity of these genes and distinguish glioma-associated expression, transcriptomic data from patients with inflammatory liver disease, specifically hepatitis[30, 31], were analysed. Analysis highlights a clear distinction between *SERPINA3* and *SPP1*. ***Fig.7(C), (D)*** While *SPP1* increases during hepatitis, *SERPINA3* does not show a similar induction. While liver is also an active producer of *SERPINA3* and *SPP1* apart from astrocytes and microglial cells, respectively, the results support the interpretation that *SERPINA3* is specific for reactive astrocytes and SPP1 seems more inflammation sensitive. In glioma, this is consistent with *SERPINA3* marking inflammatory astrocyte activation.

### Integrated co-expression, network, and enrichment analyses

Co-expression analysis is a systems-level approach that enables the identification of biological pathways activated alongside key biomarkers within a tumour. ***Fig.8(A)*** we analysed the top genes that are co-expressed with these biomarkers to gain insight into the molecular networks associated with glioma. Studying co-expressed genes helps to identify pathways and cellular processes that may be functionally linked to *SERPINA3* and *SPP1*. It reflects no neuronal genes but rather only microglial genes, meaning the microglial activation progresses the Glioma grade. Co-expression also includes extracellular matrix remodelling and invasion markers (*ANXA2, LGALS3, CD44*), and oxidative stress–metabolic adaptation genes (*SOD2, NAMPT, TXNIP*). These transcriptional signatures are characteristic of disease-associated microglia and immunosuppressive glioma microenvironments and have been repeatedly implicated in glioblastoma progression, invasion, and immune modulation. The strong correlation of these pathways with *SERPINA3* and *SPP1* supports their role as indicators of an activated, tumour-supportive microenvironment rather than isolated expression events.

**Fig 8.**
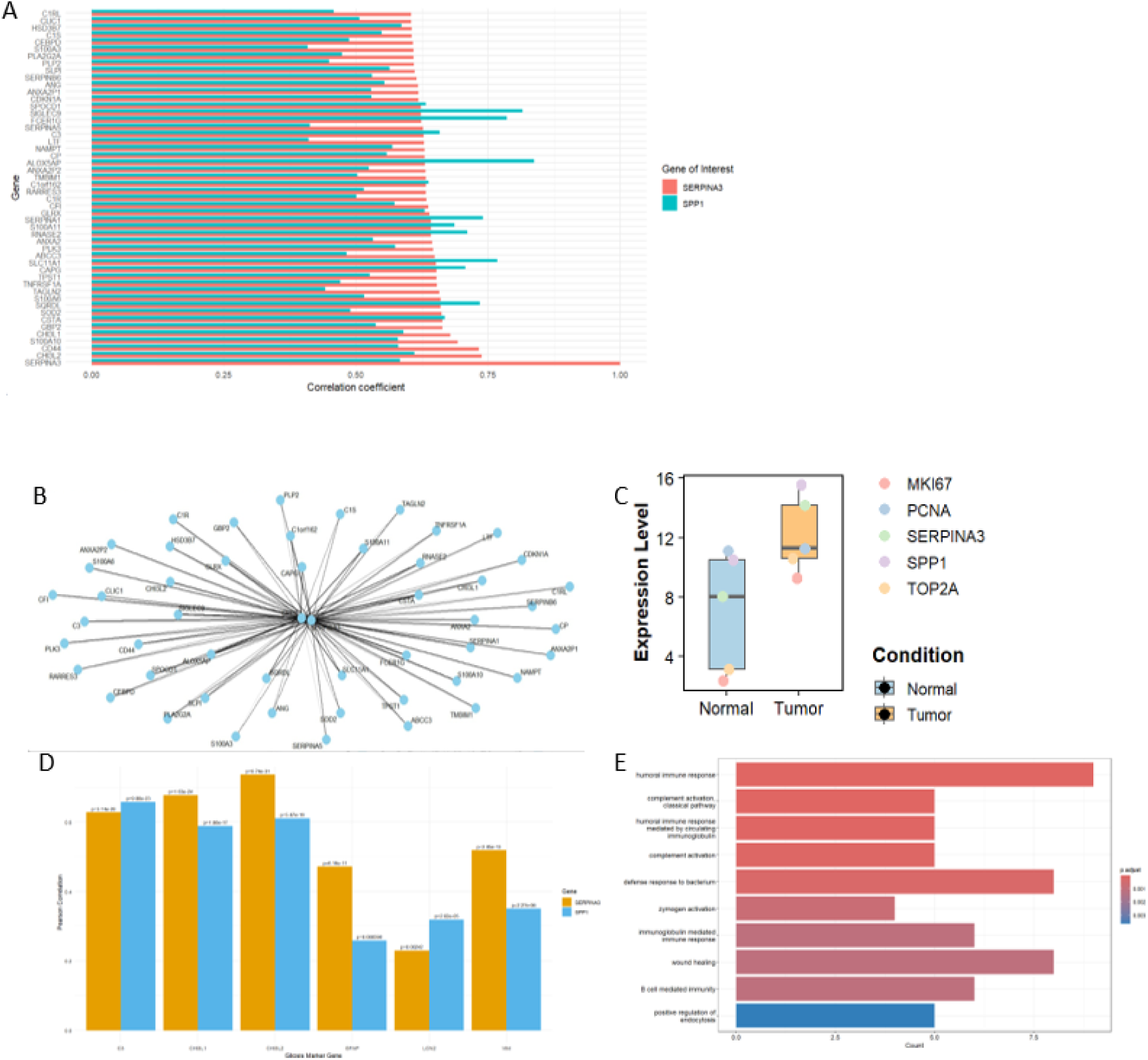
Integrated co-expression, network, and enrichment analyses **(A)** Horizontal bar plot illustrating the top 50 genes co-expressed with SERPINA3 and SPP1 ranked based on Spearman’s rank correlation coefficients. Genes were selected using statistically significant Spearman correlations (*p* < 0.05, with many correlations reaching high significance levels). **(B)** A co-expression network showcasing the top 50 genes significantly associated with *SERPINA3* and *SPP1. SERPINA3* and *SPP1* are positioned centrally, reflecting their strong connectivity with multiple co-expressed genes **(C)** Differential expression of SERPINA3 and SPP1 between normal and tumour samples. It illustrates higher expression levels of SERPINA3 and SPP1 in tumour tissues compared with normal controls. **(D)** Correlation analysis showing the association of SERPINA3 and SPP1 expression with key reactive astrocyte markers (C3, CHI3L1, CHI3L2, GFAP, LCN2, and VIM). SERPINA3 demonstrates consistently stronger correlations (*r* ≈ 0.48–0.72) with highly significant *p*-values, compared with SPP1, which shows moderate correlations (*r* ≈ 0.25–0.66). **(E)** Gene Ontology (GO) Biological Process enrichment analysis of the top 50 genes significantly co-expressed with the tumour biomarkers.

***Fig.8(B)*** Network analysis of *SERPINA3* and *SPP1* was performed where *SERPINA3* and *SPP1* are positioned centrally, reflecting their strong connectivity with multiple co-expressed genes. In ***Fig.8(C)***, to check whether *SERPINA3* and *SPP1* are linked to tumour growth, their expression was compared with proliferation markers, including *MKI67, PCNA, TOP2A, and Ki-67*. The purpose of this was to check if higher levels of *SERPINA3* and *SPP1* are associated with increased cellular proliferation in glioblastoma. The results reveal strong upregulation of *SERPINA3* and *SPP1* alongside classical proliferation markers. ***Fig.8(D)*** Correlation analysis reveals the association of SERPINA3 and SPP1 expression with key reactive astrocyte markers (C3, CHI3L1, CHI3L2, GFAP, LCN2, and VIM), supporting its role as a core astrocyte-linked inflammatory biomarker[32, 33]. Gene enrichment via gene ontology was also performed.

## Conclusion

SERPINA3 and SPP1 appeared promising progressive biomarkers in glioma, showing increased expression with tumour proliferation across LGG and GBM. The transcriptomic and proteomic data aligned with their elevated expression. Their expression by astrocytes and microglia, coupled with their involvement in inflammatory signalling and cellular crosstalk within the tumour microenvironment, suggests a functional role in glioma progression. These biomarkers cleared the specificity criteria by showing no significant association with neurodegenerative diseases like by Alzheimer’s disease, or with hepatic disorders such as hepatitis, while demonstrating glioma-specific expression in pan-cancer analyses. Furthermore, correlation analyses with markers of proliferative makers of glioma and reactive astrocytes demonstrated significant associations, suggesting that their expression arises in the context of reactive gliosis involved in inflammation. Hence, these findings indicate that the biomarkers participate in immune-mediated pathways within glioma and provide insight into the tumour microenvironment, thereby helping us to navigate through glioma progression

## Discussion

In this study, we observed increased expression of SERPINA3 and SPP1 within the glioma tumour microenvironment, particularly in regions exhibiting reactive astrogliosis, as indicated by elevated expression of astrocyte markers. Astrocytes exposed to cues from glioma cells, resident microglia, and hypoxic stress are known to acquire a reactive phenotype that has been associated with enhanced glioma proliferation, invasion, and metastatic behaviour through the release of soluble mediators and intercellular communication mediated by gap junctions. In this context, our findings extend existing models of astrocyte reactivity by identifying SPP1 and SERPINA3 as markers associated with an inflammatory and immune-modulatory axis within the glioma tumour microenvironment. The enrichment of SPP1 and SERPINA3 in tumour-associated inflammatory regions suggests that their expression reflects a broader reactive state arising from reciprocal interactions among glioma cells, astrocytes, and microglia. Notably, SERPINA3 expression was consistently higher in high-grade gliomas, supporting an association between reactive gliosis–linked inflammation and tumour aggressiveness. Furthermore, within reactive gliotic niches, glioma-derived SPP1 has been reported to interact with integrins and CD44 on tumour-associated macrophages and microglia, consistent with its association with pro-tumorigenic immune signalling in the glioma microenvironment.s

Collectively, our findings support a model where reactive astrocytes and associated glial cells promote the expression of SERPINA3 within tumour cells, simultaneously enriching the microenvironment with SPP1, which drives immune cell reprogramming and tumour support. This coordinated molecular network reinforces glioma progression. Understanding the contribution of reactive gliosis to SERPINA3 and SPP1-mediated tumour microenvironment remodelling redefines their potential as dual progressive markers that can help to keep track of the progression of the tumour and distinguish true glioma from pseudoglioma driven by reactive gliosis and inflammation. Future studies incorporating in vivo models will be essential to decipher the precise cellular sources and mechanistic roles of these molecules within the glioma microenvironment.

## Acknowledgment

**HSR, SS, SUS** acknowledge the Department of Biotechnology, Government of India for providing them the fellowship.

## CRediT authorship contribution statement

Hirtik Singh Rathore Conceptualisation, Methodology, Analysis, Investigation, Data curation, Writing – original draft, Writing – review & editing, Validation. **Siya Singh:** Investigation, Writing – review & editing. **Sukhmanpreet Singh:** Investigation. **Jassi Goyal:** Investigation, Data curation. **Dibyajyoti Banerjee:** Conceptualisation, Resources, Writing – review & editing, Supervision, Project administration.

## Data availability

Data will be made available on request.

